# Network-driven identification of indisulam neo-substrates for targeted protein degradation

**DOI:** 10.1101/2024.09.16.613231

**Authors:** Andrew F. Jarnuczak, Angelo Andres, Orli Yogev, Stephanie K. Ashenden, Cheng Ye, Fiona Pachl, Andrew Zhang, Maria Emanuela Cuomo, Meizhong Jin

## Abstract

Indisulam, a DCAF15-based molecular glue degrader, induces widespread proteome changes with implications for cell division and chromosome segregation. While RBM39 and RBM23 are two well-characterized indisulam neo-substrates, additional targets likely exist. To identify those degradation targets, we applied a network-based approach to prioritize novel neo-substrates from large-scale omics data. Our approach integrates proteome-wide expression measurements with information from publicly accessible databases into a multiplex heterogeneous network. Utilizing a Random Walk with Restart algorithm, we identified a preliminary list of 30 neo-substrates. These proteins are likely interactors with DCAF15 in the presence of indisulam and are subject to subsequent degradation. Experimental validation of RBM5, one of the shortlisted candidates, confirmed its degradation in a proteasome-dependent manner, supporting its identification as a novel indisulam neo-substrate. Our work employs established network resources and analytical methods to effectively identify direct targets of indisulam molecular glue degrader. This approach is readily adaptable for exploring novel targets across other molecular glue systems, enhancing its applicability and value to the drug discovery community.

**Summary:** Molecular glue neo-substrate identification via network analysis and proteomics profiling

## Introduction

Targeted protein degradation therapies exploit the cellular machinery to degrade disease-relevant proteins. Those therapeutics work by bringing together an E3 ubiquitin ligase (enzyme tagging proteins for degradation) and a specific target – protein of interest (POI). Bifunctional molecules, known as proteolysis-targeting chimeras (PROTACs), have two distinct binding moieties separated by a chemical linker: one for the POI and another for the E3 ligase[1]. In contrast, molecular glue degraders work by directly engaging a ubiquitin E3 ligase to form a binary complex and enable, or enhance, a ternary complex formation with the target protein.

This induced protein–protein interaction (PPI) subsequently leads to degradation of the target protein [2], [3]. Such targets are collectively termed neo-substrates.

The aryl sulfonamide compound indisulam exemplifies the molecular glue approach. Initially identified for its anti-tumour properties, it increases the affinity between an E3 ligase complex composed of DDB1 and CUL4-associated factor 15 (DCAF15) with a specific target protein, RNA-binding motif protein 39 (RBM39). This targeted interaction leads to RBM39 ubiquitination and subsequent degradation by the proteasome[4]–[6]. Interestingly, indisulam can also target the related splicing factor RBM23 through the same mechanism[7].

While RBM39 and RBM23 are two well characterized indisulam neo-substrates, there might be other yet unidentified targets, or proteins that are not *bona fide* neo-substrates, but are nevertheless affected by the compound through, for example “pathway effects”, leading to their degradation or modulation of activity. For example, it has been shown that RBM39 modulates splicing and its depletion results in up- and downregulation of transcripts and alternative splicing [8]. It is also known that indisulam disrupts cell cycle and metabolism through RBM39-mediated alternative splicing of transcripts related to these processes[9]. Most importantly, the study [9] found an overlap between proteins with reduced levels and transcripts that underwent mis-splicing upon treatment. In addition to these “pathway effects”, zinc finger E-box-binding homeobox 1 (ZEB1) has been shown to be an additional indisulam neo-substrate (indisulam degraded ZEB1 through the ubiquitin-proteasome system in a DCAF15-dependent manner) [10].

PPI data has been invaluable since the late 1990s for several applications including identification of protein multimeric complexes, elucidation of biological pathways, and prediction of protein functions. Large-scale PPI determinations have been conducted using methods such as yeast-two hybrid systems[11] or affinity chromatography paired with mass spectrometry[12]. As a result, a variety of PPI network resources have been made available publicly. Notably, primary databases that contain experimentally validated PPIs from both small-scale and large-scale studies, which are published and manually curated by experts, such as BioGRID[13] or IntAct[14].

The computational exploration of PPI networks for predicting protein complexes is also well-established as a key approach in the field. Among the numerous methodologies, the Random Walk with Restart (RWR) technique stands out due to its ability to discern multivariate relationships between nodes while capturing the global topology of the network[15]–[17]. RWR is a specialized algorithm designed to measure the closeness between two nodes within a graph. It enhances the traditional Random Walk by incorporating a restart mechanism, allowing the walker to potentially return to the starting node at each step. The algorithm iteratively updates the probabilities of being at each node by combining the probabilities from the previous step with the restart mechanism. This process continues until the probabilities converge, providing a stable distribution of likelihoods for the walker’s presence across the nodes [17], [18].

The technique has been effectively utilized to address PPI prediction or prioritization challenges. For instance, Guo et al. used quantitative proteomics data and PPI information to effectively rank proteins that are either known or suspected to be linked to drug resistance in cancer through a biased random walk model[19], while the use of RWR across various biological networks has enhanced the discovery of genetic and phenotypic associations[20].

Furthermore, novel implementations of the algorithm now accommodate its application to multiplex and heterogeneous biological networks. This extension allows RWR to navigate through various layers of physical and functional interactions among genes and proteins, including protein-protein interactions and co-expression associations. It also enables transitions across networks with differing nodes and edge types, such as those capturing phenotype similarities between diseases[17], [21]. In particular, a study describing the MultiXrank method assessed such an algorithm’s proficiency in predicting disease-associated genes, with the multiplex-heterogeneous RWR showing superior performance compared to other approaches on monoplex or heterogeneous networks[21].

Given that indisulam, and other molecular glues, must induce at the minimum transient PPIs with their targets, utilizing network or pathway-based protein association data alongside network analytics algorithms can help discover novel neo-substrates or prioritise existing information. In this work, we first characterized the broader impact of indisulam on the proteome and aimed to differentiate primary degradation effects from “pathway effects.” Leveraging results from the proteomics data, we then deployed the MultiXrank RWR-based method to prioritize potential novel indisulam-mediated neo-substrates.

## Results

### 1. Deep characterization of indisulam-induced protein expression changes

To thoroughly investigate the changes in protein expression induced by indisulam, we treated HCT116 cells with 1 µM of indisulam for either 6 or 24 hours, using six biological replicates. This approach enhanced the statistical power of our analysis, enabling us to confidently detect even smaller effect sizes.

We detected widespread modulation of the proteome with 196 downregulated and 445 upregulated proteins (minimum 20% change and adjusted p.value < 0.01) after 6h treatment. This increased after 24h (972 downregulated and 619 upregulated proteins) suggesting pathway effects are induced by indisulam treatment (Figure 1A). We note, the data revealed a wider range of changes compared to prior studies[5], [9], [10]. This likely stems from relatively high proteome coverage (quantifying 6018 proteins) and good statistical power with 6 replicates per condition.

**Figure 1.**
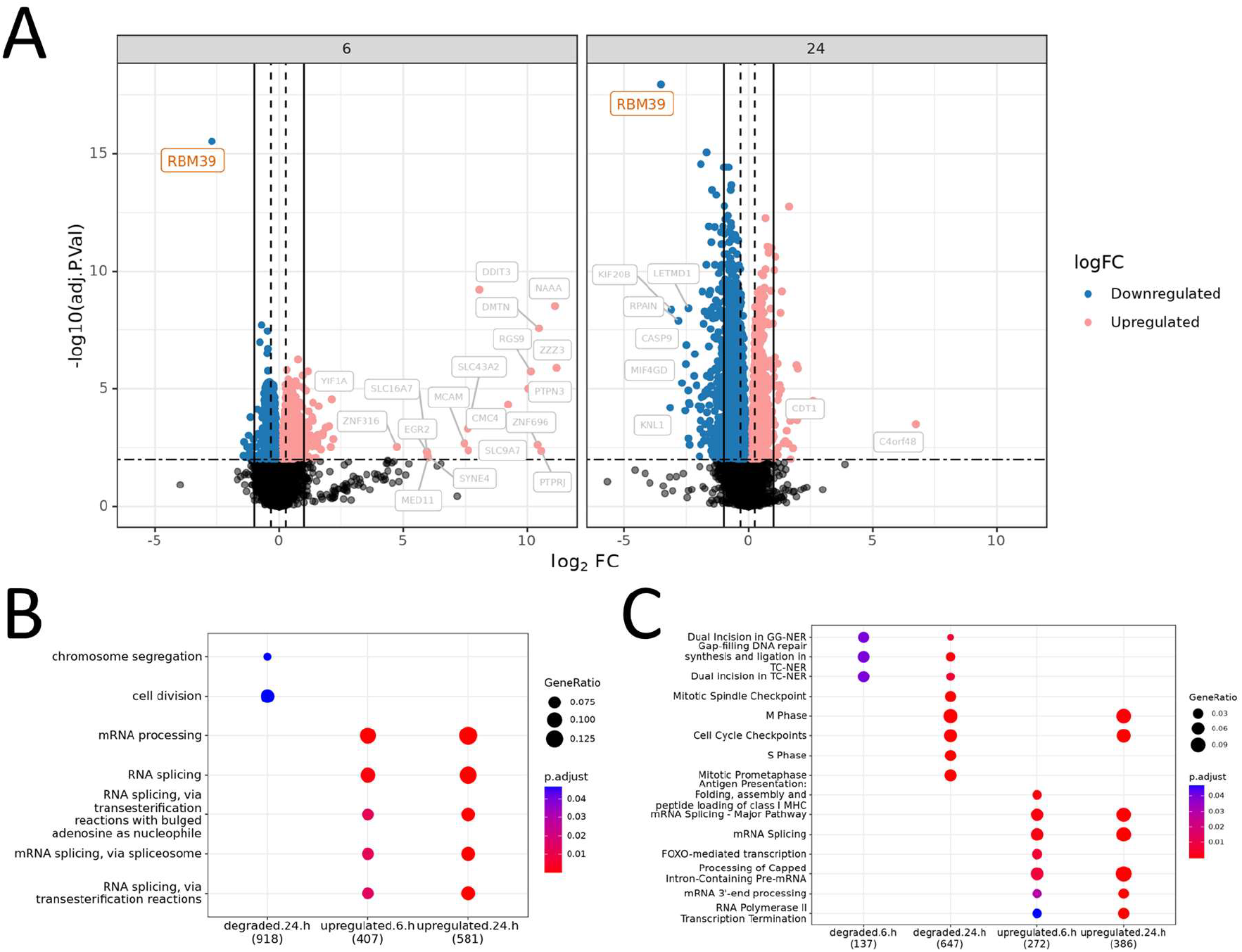
– Visualizing protein expression changes upon indisulam treatment. A) Differential expression changes after 6- and 24-hour treatment. B) Pathway Enrichment Analysis using Gene Ontology database C) Pathway Enrichment Analysis using REACTOME pathways. Note: In Figures 1B and 1C, “degraded” is used for simplicity, but this term may also encompass broader regulatory changes, including downregulation, at 6 and 24 hours.

We next utilised pathway enrichment analysis to gain deeper understanding of the targets affected directly by indisulam-induced degradation (6h degradation events) versus modulated pathways and processes (primarily proteins upregulated at both 6 and 24h). When looking at Gene Ontology (GO) biological process terms we found no enrichment among proteins regulated at 6 hours – presumably most of which would be changing due to direct degradation events. In contrast, and in agreement with previously reported studies [22], [23], we found that “cell division” and “chromosome segregation” are downregulated at 24h. Various mRNA processing GO terms are upregulated at both 6 and 24 hours (Figure 1B). In addition, analysing the Reactome pathway database[24] revealed that “M Phase” and “Cell Cycle Checkpoints” pathways were only affected at the 24h time point (Figure 1C).

This analysis allows us to begin deconvoluting direct degradation events from broader pathway effects by leveraging quantitative proteomics data.

### 2. Construction of the heterogeneous multiplex biological network

To identify which of the observed changes are specifically due to direct degradation mediated by indisulam via DCAF15, we leveraged our proteomics data alongside publicly available network databases. We hypothesized that by mining existing biological networks, we could prioritize which protein-protein interactions (PPIs) from the network(s) are most likely to occur in the presence of indisulam. These interactions, often observed under various conditions or predicted theoretically, can be effectively ranked for their likelihood of occurrence with indisulam treatment.

We constructed a multiplex heterogeneous network [25], [26], as detailed in the “Network construction and data sources” section of the Methods (Figure 2). This network comprises layers for PPIs, Gene Ontology (GO) annotations, and Reactome pathways, each layer representing distinct but interconnected datasets that describe different aspects of biological relationships and processes. The use of a heterogeneous network is particularly advantageous in this context because it allows for a structured representation of complex biological relationships across multiple dimensions and scales.

**Figure 2.**
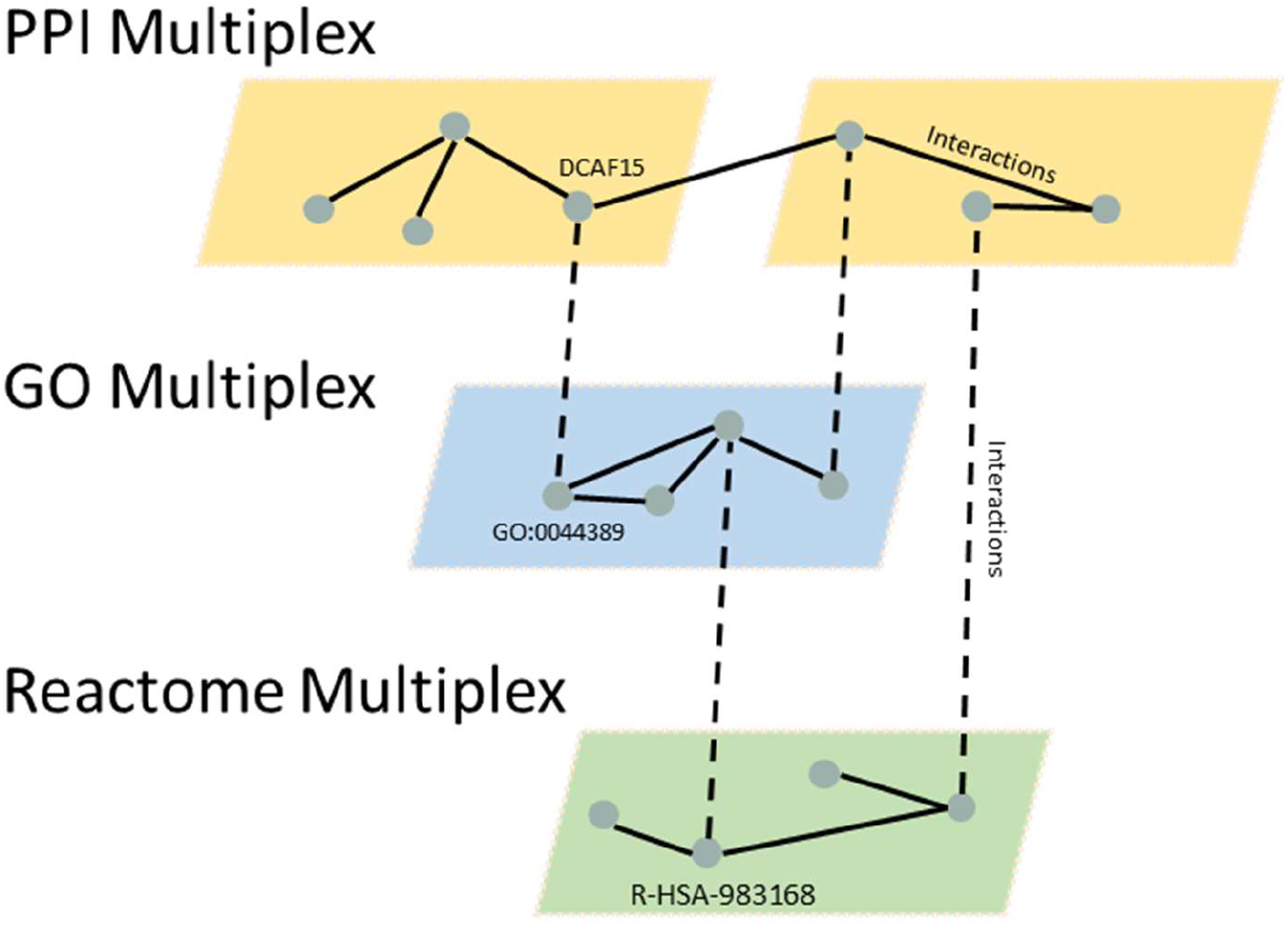
Illustration of a heterogeneous multiplex network. The heterogeneous multiplex network consists of three layers: a PPI multiplex network (upper subnetwork), GO network (middle layer) and a Reactome network (lower subnetwork). The three subnetworks are connected by the protein-GO, protein-Reactome, and GO-Reactome relationships.

### 3. Random walk with restart to prioritise indisulam-mediated DCAF15 neo substrates

We applied the RWR algorithm to our constructed network using the MultiXrank Python package[21]. RWR is a graph-traversal technique that simulates a random walker traversing the network, with a probability of returning to the current node at each step. The algorithm assigns a weight to each node, which reflects its importance within the network. The higher a node’s weight, the more frequently the RWR algorithm revisits it, indicating its central role in the network’s connectivity or “proximity” to seed nodes.

We used RWR with DCAF15 as seed or, in the final model, with multiple seeds: DDA1, DDB1, RBX1, CUL4B, CUL4A (all known to form a complex with DCAF15) and GO:0031625 (ubiquitin protein ligase binding GO term). We hypothesized that among the proteins we identified as DCAF15 neo targets for degradation, those with higher rankings in the RWR analysis are more likely to be true neo-substrates mediated by indisulam, reflecting a higher probability of interaction with DCAF15.

#### 3.1 Evaluation of RWR performance on DCAF15 dataset

To assess the performance of the method, we employed two evaluation metrics: mean reciprocal rank (MRR) and recall@k.

### Recall@k

We examined the recall performance of the ranking across different ‘k’ values for six distinct “true positive” seeds (DCAF15, DDA1, DDB1, RBX1, CUL4B, CUL4A) using the leave one out validation (LOOV) strategy. In LOOV, the RWR model is executed (trained) using all but one seed and then the model’s performance is evaluated based on that seed ranking among the top-k predicted nodes. This process is repeated for each “true positive” seed. We used a range of k from 1 (removed seed is the top ranked node) to 100 (removed seed is among top 100 ranked nodes). The results indicated the minimum “k” values to achieving 100% recall were 10 (Q16531_DDB1),18 (Q13620_CUL4B, P62877_RBX1), 22 (Q13619_CUL4A), 51 (Q66K64_DCAF15) and 60 (Q9BW61_DDA1) (Figure 3A).

**Figure 3.**
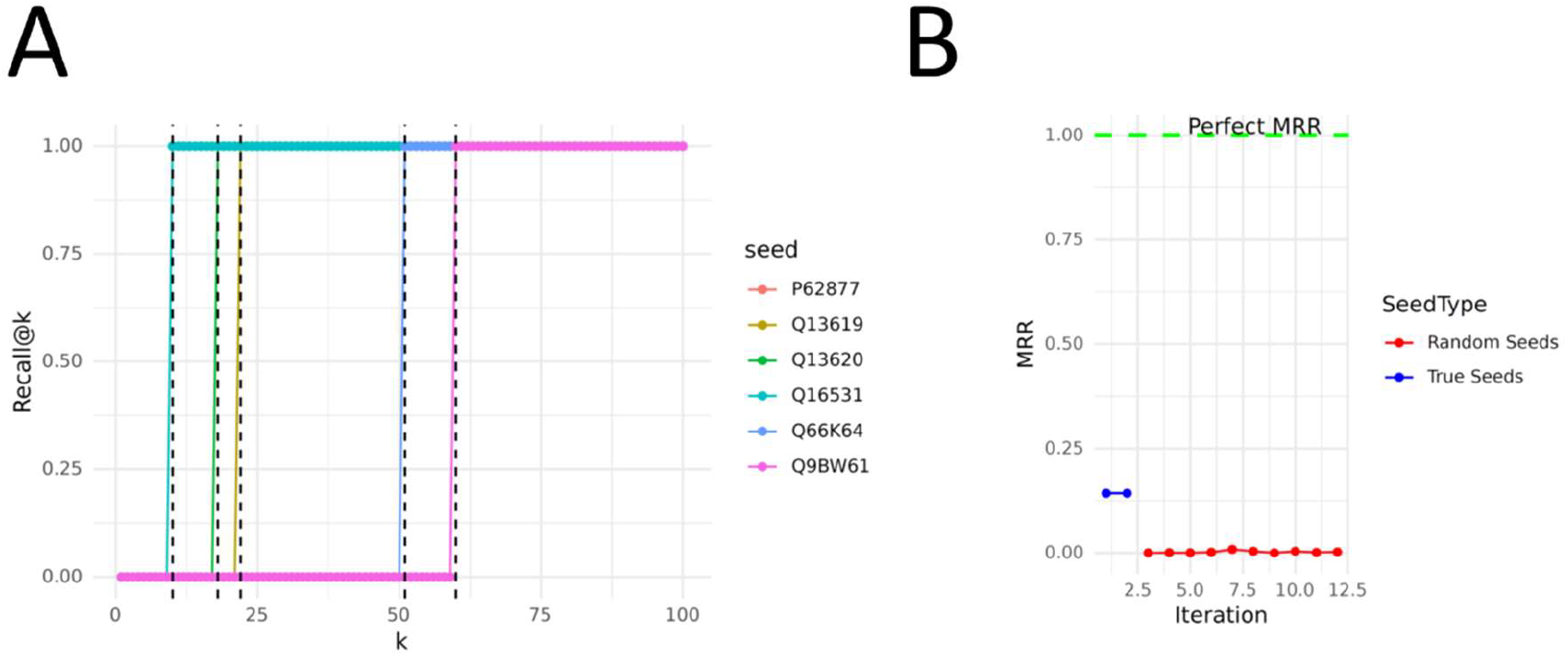
Ranking algorithm performance metrics. A) Recall performance across different ‘k’ values for each true positive seed that was removed. Dashed lines indicate the smallest ‘k’ value where 100% recall is achieved for that seed. Note, recall curves for P62877 and Q13620 seeds are overlapping. B) Comparison of MRR values for RBM39 between executing RWR ranking using True Seeds and Random Seeds. The green dashed line represents a perfect MRR score of 1. True Seeds exhibit consistently high MRR values, indicating effective ranking of RBM39 by the algorithm.

These findings indicate that selecting the top 30 ranked hits should effectively prioritize the most likely neo-substrates, despite the need for larger values of ‘k’ to ensure complete recall.

### Mean reciprocal rank (MRR)

In the MRR analysis, we assessed the ability of the algorithm to rank a single true positive indisulam-mediated neosubstrate (RBM39) using the 6 DCAF15 complex seeds (“true seeds”) versus a set of random seeds. To this end we also executed RWR for ten “random” iterations and computed the MRR metric for RBM39. MRR measures how well a ranked list of results places relevant results at the top of the list (in this case the known RBM39 network node). A high MRR score indicates that the algorithm effectively prioritized RBM39, placing it closer to the top of the list. A low MRR score suggests that the algorithm struggles to identify and position the node effectively. The results (Figure 3B) illustrate a clear distinction in the MRR values when using the “true seeds” versus random seeds. The MRR for the true seeds was 0.14, indicating a higher likelihood of the algorithm correctly identifying and ranking the true seed at the top. In contrast, the random seeds showed MRR values typically below 0.002. This distinction validates our choice of seeds and their reliability in distinguishing relevant neosubstrates from the network.

#### 3.2 Top ranked candidate DCAF15 neo-substrates

The results of the final RWR model were filtered based on the potential degradation events identified through our proteomics analysis. We retained only those hits exhibiting a minimum of 25% degradation and an adjusted p-value below 0.05 at both 6h and 24h time points. The top 30 ranked proteins from this refined set are presented in Table 1, complete with their fold changes.

**Table 1:**
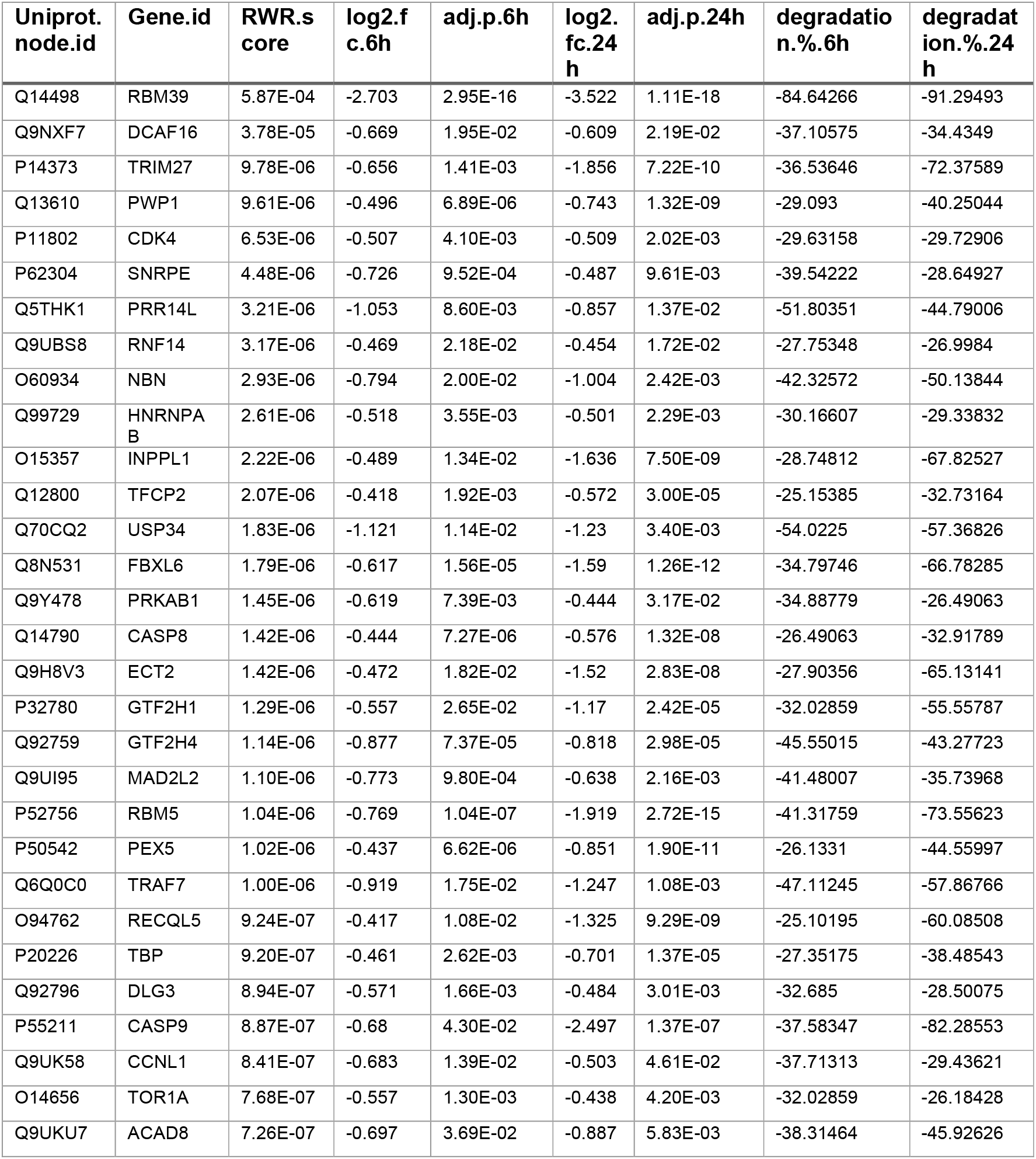
RWR scored list of top 30 proteins with significant degradation events in the proteomics assay.

The process of selecting “hits” for Table 1 involves initially ranking all candidates by their algorithm score from highest to lowest. Subsequently, any proteins not meeting the degradation criteria (25% degradation and adjusted-p value of <= 0.05 at both 6 and 24 hours from the differential expression analysis) are excluded from the list. Increasing the stringency of the degradation criteria reduces the number of hits but maintains their relative rank, focusing on those most likely affected by indisulam.

### 4. Validation of RBM5 expression changes

Among the top-ranked candidates from the RWR analysis, we selected RBM5 for experimental validation. We analysed the effect of indisulam on RBM5 protein levels using western blot on two representative cell models, HCT116 and PC9 (Figure 4A). We observed a decrease in RBM5 protein levels following indisulam treatment at 6 hours and at 24 hours, which was rescued by co-treatment with MG-132, a specific proteasome inhibitor (Figure 4B). Of interest, the levels of RBM5 degradation in HCT116 (75%) matched the proteomic data, while the levels of RBM5 degradation in PC9 cell line was to a lesser extent (40%). This finding strongly suggests that indisulam-mediated downregulation of RBM5 is dependent on the proteasome machinery and, together with network analysis, further supports its identification as a bona fide DCAF15 neo-substrate.

**Figure 4.**
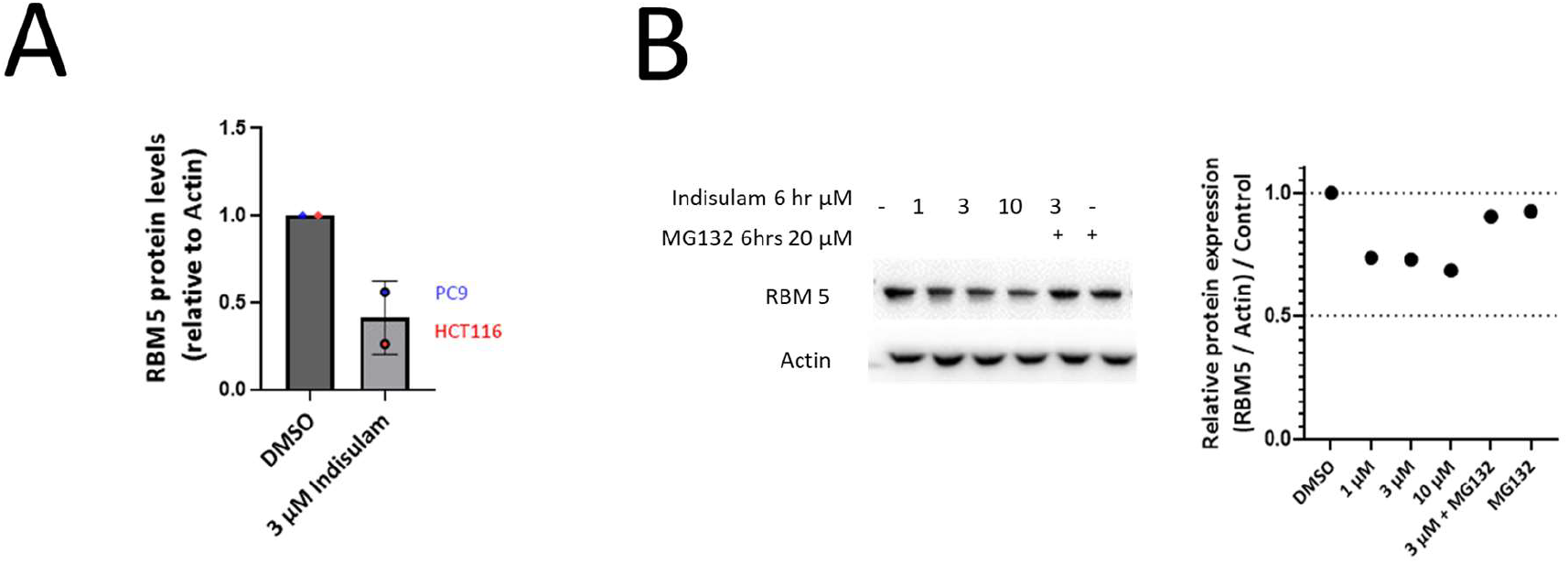
Validation of RBM5 expression changes. RBM5 protein levels following 3 µM indisulam treatment in A) PC9 (Lung cancer) and HCT116 (CRC) cell lines, 24 hrs B) HCT116 cell line, 6 hrs.

## Discussion

This study employed a combined approach of RWR network analysis and proteomics to identify indisulam-mediated neo-substrates of DCAF15. Our findings reveal a set of downregulated proteins following indisulam treatment, with several strong candidates emerging from the RWR analysis. Notably, the top-ranked candidate, RBM39, aligns with previous reports[4], [9], [23].

The proteomics data revealed a large number of proteome changes compared to some previous studies employing proteomics following indisulam treatment [10]. This is due to two key factors. Firstly, we achieved high proteome coverage, quantifying a significant number of proteins (6018 proteins) across two treatment time-points. This comprehensive analysis provides a broader picture of indisulam effects. Secondly, our experimental design prioritized statistical power by incorporating six biological replicates.

Our pathway enrichment analysis differentiates between proteins downregulated at 6h (likely enriched with direct degradation neo-substrates) and those showing changes at both 6h and 24h (this group may represent a combination of direct degradation and modulation of pathways influenced by the treatment). This distinction is crucial because initial degradation should occur relatively fast. Proteins downregulated at 6h likely represent direct interactions with the drug. Ubiquitination and subsequent proteasomal degradation typically occur within a few hours[27], [28]. Observing degradation at this early time point strengthens the evidence for a direct neo-substrate relationship. In contrast downstream effects (indirect consequences of indisulam activity) are likely to happen later. For example, disruption of a key pathway components by indisulam at 6h can lead to downstream effects, including the degradation of additional proteins involved in the same pathway later point (24h).

Some limitations of this interpretation include turnover rates and indirect effects at 6h. Turnover Rates: While early degradation is suggestive, it’s important to acknowledge that protein turnover rates can vary[29]–[31]. Therefore, some neo-substrates might have slower degradation kinetics, leading to a delay in their disappearance. Indirect Effects at 6h: In some cases, even proteins degraded at 6h could be indirectly affected by indisulam. However, the earlier time point increases the likelihood of a direct interaction.

We also sought to reference previously published DCAF15 interactome data generated upon indisulam treatment[32], [33]. While some overlap was observed (e.g., RBM39), there was a limited number of shared hits (out of 134 significant hits from our data only 9 were detected in[33]).

Interestingly, our data revealed significant degradation of JMJD1C following indisulam treatment; however, JMJD1C was not ranked highly in our network analysis. This finding aligns well with the literature, where JMJD1C was detectable but not highlighted as a prominent DCAF15 interactor [33], This suggests that our network analysis accurately reflects the literature’s characterization of JMJD1C’s interaction with DCAF15.

## Methods

### Global Proteomics Sample Preparation

HCT116 cells grown in McCoy’s 5A growth medium (supplemented with 10% FBS and 1% L-Glutamine) were seeded into 6-well tissue culture plates at 700,000 cells/mL in 1 mL. After overnight incubation at 37° C in a 5% CO2 atmosphere, growth medium was replaced with growth media treated with 0.1% DMSO or 1 μM indisulam. After 6 hour and 24-hour incubation the growth medium was removed, cells were washed with 1 mL PBS, and cells were resuspended using 0.5 mL 0.5% trypsin solution for 5 min at 37°C. Trypsin was neutralized using equal volume of growth medium and cells were then transferred into 1.5 mL Eppendorf tubes for centrifugation (700g x 2 min) and three wash cycles using ice-cold PBS. After removing the final PBS wash the cell pellets were snap-frozen over dry ice and stored at - 80°C until lysis.

HCT116 cell extracts were prepared using iST 96x sample preparation kit according to manufacturer’s instructions (PreOmics). In short, cell pellets were lysed by resuspension in a lysis buffer solution and heated to 95 °C for 10 min shaking (1000 rpm) followed by pulsed sonication (10x 1sec pulses, Qsonica). Aliquots containing 50 µg of protein were transferred into cartridges, digestion solution was added and proteins were digested for 3 hours at 37 °C. Digestion was stopped by adding “stop” solution and peptides were purified by centrifugation for 2 min at 2000 g, followed by two washes and elution into a collection plate using provided solutions. Peptides were dried in a vacuum centrifuge and resuspended in 0.1% formic acid, 2% acetonitrile (ACN) for MS analysis.

### LC-MS/MS Analysis

Nanoflow LC-MS/MS was performed by coupling an Ultimate 3000 RSLCnano system to an Orbitrap Eclipse mass spectrometer. 2.2 µg peptides were trapped on an Acclaim Pepmax column (2 cm x 75 µm, 3 µm, 100 Å) in water with 0.1% formic acid (FA) and separated on a C18 Dr. Maisch column (24 cm x 75µm, 3 µm) at 50 °C. Gradient elution was performed from 2.7% ACN to 27% ACN in 0.1% FA over 118 min at 250 nL/min flow rate. Mass spectrometer was operated in data-independent acquisition (DIA) mode with full scan MS spectra acquired at 120,000 resolution and AGC target at 100%. MS2 spectra were collected at 15,000 resolution, AGC target at 100% and normalized collision energy set at 28%. 75 variable windows covering a mass range of 400-1000 m/Z and with an overlap of 1 m/z were used.

### RBM5 Validation Assay

The cell lines HCT116 (ATCC) and PC9 (ECACC), which were STR validated and tested negative for mycoplasma, were cultured in RPMI 1640 media with GlutaMAX supplement and 9% FBS at 37°C in a 5% CO2 atmosphere. The cells were treated 24 hours after plating with 3µM Indisulam (SML1225-5MG, Sigma) with or without 20µM MG132 (M7449-200UL, Sigma) for either 24 hours or 6 hours. Following the treatment, the cells were lysed using RIPA buffer (89901, Thermo Scientific) supplemented with protease and phosphatase inhibitors. Samples were run on 4-12% SDS-PAGE gels and separated by electrophoresis. The proteins were then transferred using iBlot2 (Invitrogen) and blocked with 5% non-fat milk for 1 hour at room temperature. Subsequently, the membrane was probed with primary antibodies against RBM5 (HPA018011, Sigma) and beta-Actin (3700S, Cell Signalling Technology) overnight at 4°C, followed by incubation with secondary antibodies (IRDye800 Goat anti-rabbit IgG, 926-32211, and IRDye680 Donkey anti-mouse IgG, 926-68022, Li-COR). Protein bands were visualized using ODYSSEY CLx and quantified using ImageJ software.

### Mass Spectrometry Data Processing

Raw files were searched with Pulsar embedded in Spectronaut (version 17.3) against a UniProt human database (February 2021) using directDIA workflow and default BGS factory settings with Trypsin/P digestion and two missed cleavages allowed. Cysteine carbamidomethylation was set as a fixed modification, while methionine oxidation and N-terminal acetylation were set as variable modifications. Cross-run normalization was performed within Spectronaut.

### Differential Protein Expression Analysis

Normalized protein abundance values obtained from Spectronaut were evaluated for possible batch effects and sample quality utilizing principal component analysis and unsupervised hierarchical clustering. An initial filter retained proteins quantified in at least 60% of the samples before performing differential expression analysis. The analysis was done in R (version 4.1.1) using Empirical Bayes statistics within protein-wise linear models, facilitated by the limma package [34], and incorporated in the in-house data analytics portal. Proteins significantly down/upregulated in the treatment samples compared to DMSO controls were identified with a threshold of adjusted p.value < 0.01 and a minimum 20% change (where % Change=(2^^log2(fold change)^ −1)×100).

### Network Construction and Data Sources

In a homogeneous network, all nodes, and interactions, regardless of their domain-specific properties, are treated equally. Conversely, a heterogeneous multiplex network, as depicted in Figure 2, organizes different types of data into distinct layers. Each layer focuses on a specific type of biological entity or interaction.

In our network the PPI sub-network contained four layers: data from the Biological General Repository for Interaction Datasets (BioGRID)[13] (file ppi.biogrid.uniprot.mapped.2023.12.05.tsv), database of “E3-Substrate interactions” (file ubinet-E3_substrate_interactions.tsv downloaded from https://awi.cuhk.edu.cn/~ubinet/download.php) and “Protein - Protein interactions” (file httpsawi.cuhk.edu.cn.ubinetdownload.php_PPI_Human_210209.tsv downloaded from https://awi.cuhk.edu.cn/~ubinet/download.php) from UbiNet 2.0[35] and predicted E3-substrate interactions from the Degpred resource[36] (file httpdegron.phasep.pro_predicted_ESIs.tsv downloaded from http://degron.phasep.pro/download/). The GO network layer was constructed using gene ontology database which is a structured and standardized vocabulary used to describe gene functions, processes, and locations[37]. In this network, nodes represented GO terms and edges the relationships between the nodes. A GO term-protein mapping is connecting proteins to their GO classification. Similarly, Reactome pathways hierarchy was converted to the third network layer. The final graph contained intra-network edges between nodes of the same type (e.g. GO-GO term edges) and inter-network connections linking nodes of different types (e.g. GO term – protein edge) (Figure 2).

### Prioritizing Degradation Events via Network Proximity Ranking with RWR

To identify which degradation events in our experimental dataset are likely due to direct interactions with the seed nodes (such as DCAF15 and RBM39), we followed a systematic process involving 3 steps:

1. Execution of the Random Walk with Restart (RWR) Algorithm: We first ran the RWR algorithm on our network using the MultiXrank Python package[21], a comprehensive framework specifically designed for executing RWR on universal multilayer networks. MultiXrank takes several parameters, including a configuration file and seed nodes, to tailor the analysis to the specific dataset. The configuration file details the structure of the multilayer network and specifies the bipartite connections between different layers. The seeds used here were: GO:0031625, Q66K64, Q9BW61, Q16531, P62877, Q13620, Q13619
2. Generation of Network Proximity Rankings: The output of the RWR process is a ranking of all proteins, reflecting their network proximity to seed nodes.
3. Integration with Experimental Expression Data: We then refined the initial proximity rankings by integrating them with fold change data from our proteomics analysis. Specifically, we filtered the ranked list to retain only those proteins that showed significant degradation, defined as at least 25% degradation with an adjusted p-value below 0.05 at both the 6-hour and 24-hour time points. From this refined set, the top 30 proteins are presented in Table 1.

## Data and Code Availability

The data and software used in this study are publicly available. The script and configuration files to execute MultiXrank have been deposited in a GitHub repository, which can be accessed here https://github.com/J-Andy/molecular-glue-network-prediction.

Additional data generated at various stages of the computational processes are available upon request.

## Acknowledgements

We extend our sincere thanks to Edward Morrissey for his insightful discussions and providing expertise on best practices in knowledge graph and network analysis.

## References

[1] M. Békés, D. R. Langley, and C. M. Crews, “PROTAC targeted protein degraders: the past is prologue,” Nat. Rev. Drug Discov. 2022 213, vol. 21, no. 3, pp. 181–200, Jan. 2022, doi: 10.1038/s41573-021-00371-6.

[2] J. M. Kastl, G. Davies, E. Godsman, and G. A. Holdgate, “Small-Molecule Degraders beyond PROTACs—Challenges and Opportunities,” SLAS Discov., vol. 26, no. 4, pp. 524–533, Apr. 2021, doi: 10.1177/2472555221991104.

[3] Y. Che, A. M. Gilbert, V. Shanmugasundaram, and M. C. Noe, “Inducing protein-protein interactions with molecular glues,” Bioorg. Med. Chem. Lett., vol. 28, no. 15, pp. 2585–2592, Aug. 2018, doi: 10.1016/J.BMCL.2018.04.046.

[4] T. Han et al., “Anticancer sulfonamides target splicing by inducing RBM39 degradation via recruitment to DCAF15,” Science (80-.)., vol. 356, no. 6336, Apr. 2017, doi: 10.1126/SCIENCE.AAL3755/SUPPL_FILE/AAL3755-HAN-SM.PDF.

[5] T. B. Faust et al., “Structural complementarity facilitates E7820-mediated degradation of RBM39 by DCAF15,” Nat. Chem. Biol. 2019 161, vol. 16, no. 1, pp. 7–14, Nov. 2019, doi: 10.1038/s41589-019-0378-3.

[6] T. Uehara et al., “Selective degradation of splicing factor CAPERα by anticancer sulfonamides,” Nat. Chem. Biol. 2017 136, vol. 13, no. 6, pp. 675–680, Apr. 2017, doi: 10.1038/nchembio.2363.

[7] T. C. Ting et al., “Aryl Sulfonamides Degrade RBM39 and RBM23 by Recruitment to CRL4DCAF15,” Cell Rep., vol. 29, no. 6, pp. 1499-1510.e6, Nov. 2019, doi: 10.1016/J.CELREP.2019.09.079.

[8] S. Campagne et al., “Molecular basis of RNA-binding and autoregulation by the cancer-associated splicing factor RBM39,” Nat. Commun. 2023 141, vol. 14, no. 1, pp. 1–18, Sep. 2023, doi: 10.1038/s41467-023-40254-5.

[9] A. Nijhuis et al., “Indisulam targets RNA splicing and metabolism to serve as a therapeutic strategy for high-risk neuroblastoma,” Nat. Commun. 2022 131, vol. 13, no. 1, pp. 1–16, Mar. 2022, doi: 10.1038/s41467-022-28907-3.

[10] J. Lu et al., “The aryl sulfonamide indisulam inhibits gastric cancer cell migration by promoting the ubiquitination and degradation of the transcription factor ZEB1,” J. Biol. Chem., vol. 299, no. 4, Apr. 2023, doi: 10.1016/j.jbc.2023.103025.

[11] S. Fields and R. Sternglanz, “The two-hybrid system: an assay for protein-protein interactions,” Trends Genet., vol. 10, no. 8, pp. 286–292, Aug. 1994, doi: 10.1016/0168-9525(90)90012-U.

[12] J. H. Morris et al., “Affinity purification–mass spectrometry and network analysis to understand protein-protein interactions,” Nat. Protoc. 2014 911, vol. 9, no. 11, pp. 2539–2554, Oct. 2014, doi: 10.1038/nprot.2014.164.

[13] R. Oughtred et al., “The BioGRID database: A comprehensive biomedical resource of curated protein, genetic, and chemical interactions,” Protein Sci., vol. 30, no. 1, p. 187, Jan. 2021, doi: 10.1002/PRO.3978.

[14] N. del Toro et al., “The IntAct database: efficient access to fine-grained molecular interaction data,” Nucleic Acids Res., vol. 50, no. D1, pp. D648–D653, Jan. 2022, doi: 10.1093/NAR/GKAB1006.

[15] M. Rosvall and C. T. Bergstrom, “Maps of random walks on complex networks reveal community structure,” Proc. Natl. Acad. Sci. U. S. A., vol. 105, no. 4, pp. 1118–1123, Jan. 2008, doi: 10.1073/PNAS.0706851105/SUPPL_FILE/06851SUPPAPPENDIX.PDF.

[16] J. Peng et al., “Improving the measurement of semantic similarity by combining gene ontology and co-functional network: a random walk based approach,” BMC Syst. Biol., vol. 12, no. Suppl 2, Mar. 2018, doi: 10.1186/S12918-018-0539-0.

[17] A. Valdeolivas et al., “Random walk with restart on multiplex and heterogeneous biological networks,” Bioinformatics, vol. 35, no. 3, pp. 497–505, Feb. 2019, doi: 10.1093/BIOINFORMATICS/BTY637.

[18] F. Xia, J. Liu, H. Nie, Y. Fu, L. Wan, and X. Kong, “Random Walks: A Review of Algorithms and Applications,” IEEE Trans. Emerg. Top. Comput. Intell., vol. 4, no. 2, pp. 95–107, Apr. 2020, doi: 10.1109/TETCI.2019.2952908.

[19] H. Guo et al., “Biased random walk model for the prioritization of drug resistance associated proteins,” Sci. Reports 2015 51, vol. 5, no. 1, pp. 1–14, Jun. 2015, doi: 10.1038/srep10857.

[20] Y. Li and J. C. Patra, “Genome-wide inferring gene–phenotype relationship by walking on the heterogeneous network,” Bioinformatics, vol. 26, no. 9, pp. 1219–1224, May 2010, doi: 10.1093/BIOINFORMATICS/BTQ108.

[21] A. Baptista, A. Gonzalez, and A. Baudot, “Universal multilayer network exploration by random walk with restart,” Commun. Phys. 2022 51, vol. 5, no. 1, pp. 1–9, Jul. 2022, doi: 10.1038/s42005-022-00937-9.

[22] T. Owa et al., “Discovery of novel antitumor sulfonamides targeting G1 phase of the cell cycle,” J. Med. Chem., vol. 42, no. 19, pp. 3789–3799, Sep. 1999, doi: 10.1021/JM9902638/ASSET/IMAGES/MEDIUM/JM9902638U00003A.GIF.

[23] D. E. Bussiere et al., “Structural basis of indisulam-mediated RBM39 recruitment to DCAF15 E3 ligase complex,” Nat. Chem. Biol. 2019 161, vol. 16, no. 1, pp. 15–23, Dec. 2019, doi: 10.1038/s41589-019-0411-6.

[24] M. Milacic et al., “The Reactome Pathway Knowledgebase 2024,” Nucleic Acids Res., vol. 52, no. D1, pp. D672–D678, Jan. 2024, doi: 10.1093/NAR/GKAD1025.

[25] B. Lee, S. Zhang, A. Poleksic, and L. Xie, “Heterogeneous Multi-Layered Network Model for Omics Data Integration and Analysis,” Front. Genet., vol. 10, p. 501269, Jan. 2020, doi: 10.3389/FGENE.2019.01381/BIBTEX.

[26] M. De Domenico et al., “Mathematical Formulation of Multi-Layer Networks,” Phys. Rev. X, vol. 3, no. 4, Jul. 2013, doi: 10.1103/PhysRevX.3.041022.

[27] K. M. Riching, E. A. Caine, M. Urh, and D. L. Daniels, “The importance of cellular degradation kinetics for understanding mechanisms in targeted protein degradation,” Chem. Soc. Rev., vol. 51, no. 14, pp. 6210–6221, Jul. 2022, doi: 10.1039/D2CS00339B.

[28] E. F. Vieux et al., “A Method for Determining the Kinetics of Small-Molecule-Induced Ubiquitination,” SLAS Discov., vol. 26, no. 4, pp. 547–559, Apr. 2021, doi: 10.1177/24725552211000673.

[29] A. B. Ross, J. D. Langer, and M. Jovanovic, “Proteome Turnover in the Spotlight: Approaches, Applications, and Perspectives,” Mol. Cell. Proteomics, vol. 20, p. 100016, 2021, doi: 10.1074/MCP.R120.002190.

[30] Z. Rolfs, B. L. Frey, X. Shi, Y. Kawai, L. M. Smith, and N. V. Welham, “An atlas of protein turnover rates in mouse tissues,” Nat. Commun. 2021 121, vol. 12, no. 1, pp. 1–9, Nov. 2021, doi: 10.1038/s41467-021-26842-3.

[31] E. F. Fornasiero and J. N. Savas, “Determining and interpreting protein lifetimes in mammalian tissues,” Trends Biochem. Sci., vol. 48, no. 2, pp. 106–118, Feb. 2023, doi: 10.1016/J.TIBS.2022.08.011.

[32] S. Yamanaka et al., “A proximity biotinylation-based approach to identify protein-E3 ligase interactions induced by PROTACs and molecular glues,” Nat. Commun. 2022 131, vol. 13, no. 1, pp. 1–17, Jan. 2022, doi: 10.1038/s41467-021-27818-z.

[33] J. Lu et al., “Proximity Labeling, Quantitative Proteomics, and Biochemical Studies Revealed the Molecular Mechanism for the Inhibitory Effect of Indisulam on the Proliferation of Gastric Cancer Cells,” J. Proteome Res., vol. 20, no. 9, pp. 4462–4474, Sep. 2021, doi: 10.1021/ACS.JPROTEOME.1C00437/ASSET/IMAGES/LARGE/PR1C00437_0008.JPEG.

[34] M. E. Ritchie et al., “limma powers differential expression analyses for RNA-sequencing and microarray studies,” Nucleic Acids Res., vol. 43, no. 7, pp. e47–e47, Apr. 2015, doi: 10.1093/NAR/GKV007.

[35] Z. Li et al., “UbiNet 2.0: a verified, classified, annotated and updated database of E3 ubiquitin ligase–substrate interactions,” Database, vol. 2021, Sep. 2021, doi: 10.1093/DATABASE/BAAB010.

[36] C. Hou, Y. Li, M. Wang, H. Wu, and T. Li, “Systematic prediction of degrons and E3 ubiquitin ligase binding via deep learning,” BMC Biol., vol. 20, no. 1, pp. 1–19, Dec. 2022, doi: 10.1186/S12915-022-01364-6/FIGURES/7.

[37] T. G. O. Consortium et al., “The Gene Ontology knowledgebase in 2023,” Genetics, vol. 224, no. 1, May 2023, doi: 10.1093/GENETICS/IYAD031.

